# A novel enterococcus phage endolysin Lys22 with a wide host range against mixed biofilm of *Enterococcus faecalis*, *Staphylococcus aureus* and *Acinetobacter baumannii*

**DOI:** 10.1101/2025.03.11.642608

**Authors:** Ziqin Yang, Xue Du, Nannan Hu, Meng-ai Feng, Jiaoyang Xu, Hailin Jiang, Na Zhang, Honglan Huang, Jinghua Li, Hongyan Shi

## Abstract

The global surge in multidrug-resistant (MDR) bacterial pathogens has created an urgent imperative for innovative antimicrobial strategies. *Enterococcus faecalis*, *Staphylococcus aureus*, and *Acinetobacter baumannii* demonstrate remarkable antibiotic resistance and dominate hospital-acquired infections. These bacteria often form biofilms, a complex community structure that shields them from immune system phagocytosis, resists antibiotic penetration, and enhances their survival in harsh environments. In clinical cases, these bacteria often form mixed biofilms and lead to treatment failures. Phages and their derivatives have emerged as promising candidates in the fight against drug-resistant bacteria. Lys22, an endolysin derived from an enterococcus phage, has been cloned and demonstrated to possess a broad host range, effectively targeting *E. faecalis*, various *Staphylococcus* species, and *A. baumannii*. When applied to the biofilms formed by these bacteria, Lys22 was found to significantly inhibit both simple and complex biofilms in vitro. Virulent genes, including *agrA, sarA*, and *ic*aA in *S. aureus*; *asa1*, *cylA*, and *hyl* in *E. faecalis*; and *OmpA* and *lpsB* in *A. baumannii* were also downregulated by Lys22. Notably, Lys22 also exhibited a robust protective effect against dual or triple infections involving *E. faecalis*, *S. aureus*, and *A. baumannii* in a zebrafish eggs model, highlighting its potential as a therapeutic agent in combatting multi-bacterial infections.

## Introduction

*Enterococci, Staphylococcus aureus,* and *Acinetobacter baumannii* are priority members of resistant bacteria that need to explore effective new antimicrobial agents ^[1]^*. Enterococci* are very important pathogens of nosocomial infections, and *E. faecalis* occupies about 80% of enterococcal clinical isolates ^[2]^. *E. faecalis* showed strong tolerance to some extreme environments including low pH, high concentration of NaCl, poor nutrition and some antibiotics. *E. faecalis* has intrinsic and acquired resistance to many antibiotics through encoding low affinity antibiotic binding protein, efflux pumps, low membrane permeability, modifying enzymes, ribosome protection proteins and changes of antibiotic targets ^[1, 3]^. Notably, biofilm formation serves as a multifunctional survival strategy of *E. faecalis*, enabling antibiotic evasion, facilitating bacterial dissemination, and acting as a reservoir for resistance gene exchange ^[4, 5]^.

Similarly, *S. aureus* remains a preeminent global pathogen causing pneumonia, prosthetic joint infections, surgical, surgical site complications, and healthcare-associated bacteriemia ^[6]^. The emergence of pan-drug-resistant strains has elevated this pathogen to a critical priority in antimicrobial development ^[7]^. Like *E. faecalis*, its capacity to form tenacious biofilms on both biotic and abiotic surfaces significantly contributes to therapeutic failure in clinical settings ^[8-10]^.

Among Gram-negative pathogens, *A. baumannii* stands out as a notorious cause of hospital-acquired infections, with its innate multidrug resistance frequently rendering conventional therapies ineffective ^[11]^. Especially, carbapenem-resistant strains occupied first place of WHO’s critical priority list for antibiotic development ^[12]^. The pathogen’s extraordinary environmental persistence stems from its remarkable adaptive mechanisms, particularly through biofilm-mediated colonization of both living tissues and medical devices, thereby establishing protected reservoirs against antimicrobial agents ^[13-15]^.

Clinically relevant biofilms rarely exist as mono-species communities, with persistent infections frequently involving complex polymicrobial consortia ^[16]^. The frequent coexistence of *E. faecalis* and A. baumannii with S. aureus in chronic infections exacerbates clinical outcomes, as mixed biofilm not only prolong patient recovery but also accelerates the evolution of drug-resistant variants ^[17]^.

Phage-derived endolysins, which lyse bacterial cells by hydrolyzing peptidoglycan bonds during progeny phage release, have shown promise in biofilm eradication ^[18]^. However, their narrow specificity limits their application against polymicrobial infections ^[19,20]^.

In this study, we characterized Lys22, a novel endolysin originated from Enterococcus phage LY0322. This enzyme demonstrated broad-spectrum activity against *E. faecalis*, multiple *Staphylococcus* species, *A. baumannii*, *A. pittii*, *A. nosocomialis*, and *Enterobacter hormaechei*. We subsequently evaluated its anti-biofilm efficacy against single-, dual-, and triple-pathogen consortia involved E. faecalis, S. aureus, and A. baumannii in vitro, and assessed its protective effects in zebrafish embryo infection models challenged with these pathogens.

## Result

### Clone and expression of Lys22

The endolysin from phage LY0322 is a N-acetylmuramoyl-L-alanine amidase compromising 331 amino acids (Protein ID: YP_009624708.1). Structurally, the N-terminal region (residues 46-211) contains a conserved CwIA domain characteristic of peptidoglycan hydrolases that maintain the equilibrium between cell wall synthesis and hydrolysis in Gram-positive bacteria, as documented in *C. difficile* and *Bacillus thuringiensis* ^[21,22]^. Critical zinic-binding residues (His52, His160, and Asp173) form a catalytic triad within the three-dimension structure (Fig.1). The C-terminal ZoocinA-TRD (target recognition domain) features a β-sheet architecture with antiparallel strands and a short α-helix, mediating species-specific substrate binding.

**Fig. 1.**
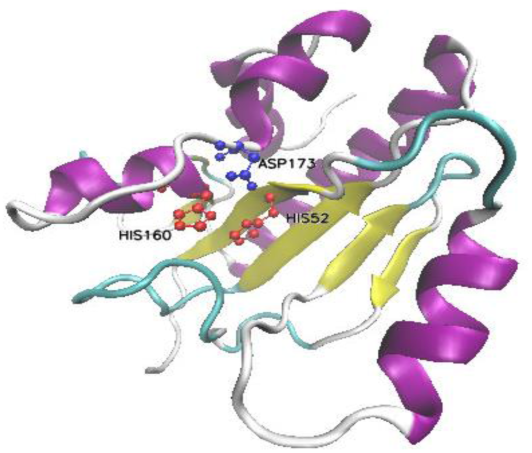
Structure of Lys22. 3D construction of Lys22, which was performed in online SWISS-MODEL server.

### Host range of endolysin Lys22

Primarily, the host range of Lys22 was investigated in Gram positive bacteria, 36 strains of *E. faecalis* and 49 strains of staphylococci were used as target bacteria in dropping lawn tests. Lys22 demonstrated broad lytic activity to 11 strains of *E. faecalis* and 24 strains of staphylococci. The sensitive staphylococci include *S. aureus* (10), *S. epidermidis* (4), *S. haemolyticus* (5), *S. cohnii* (1), *S. hominis* (1), *S. klosii* (1) and *S. warneri* (1) (Table 1).

**Table 1.**
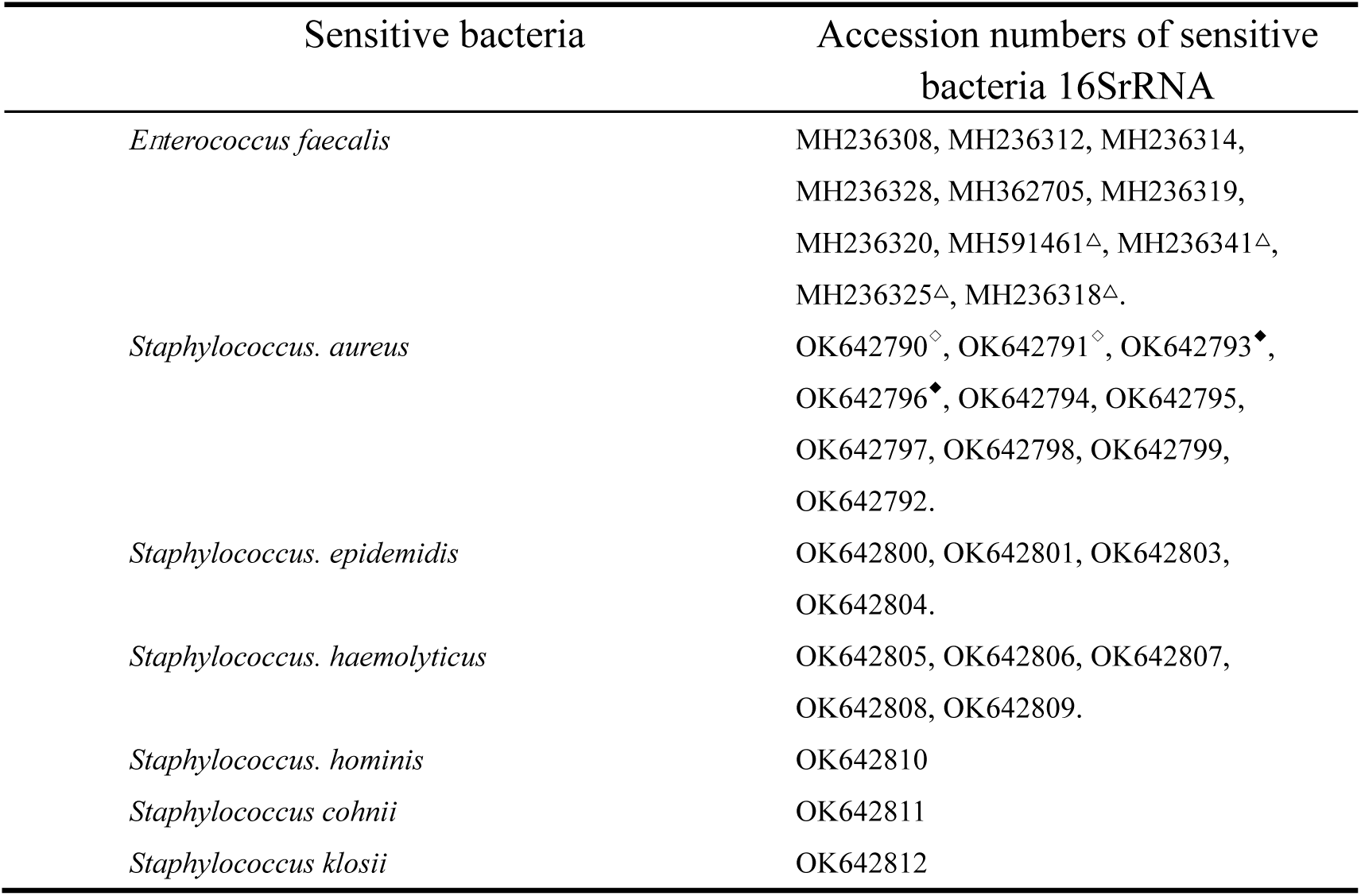
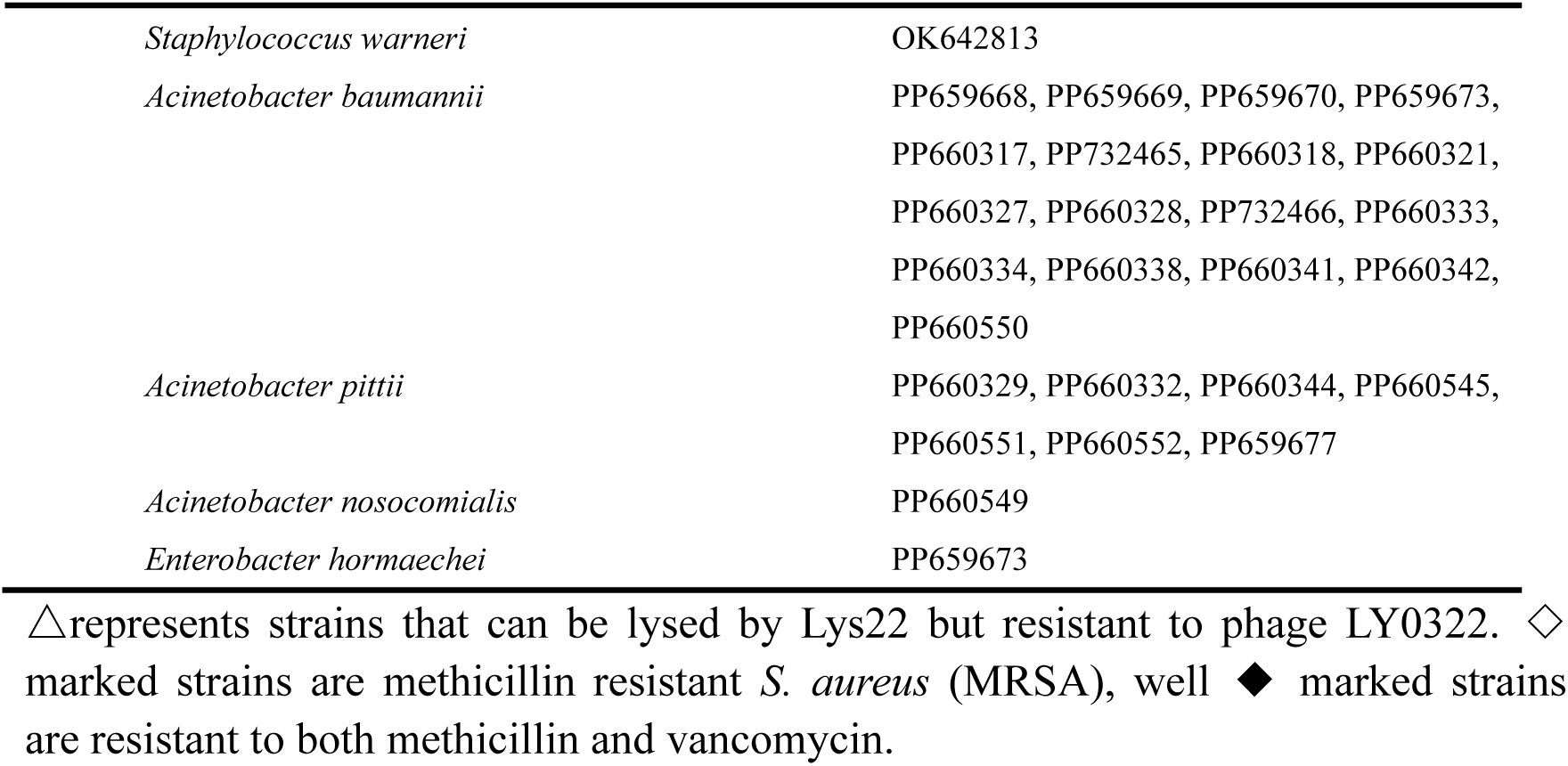
Bacteria lysed by Lys22.

| Sensitive bacteria | Accession numbers of sensitive bacteria 16SrRNA |
| --- | --- |
| *Enterococcus faecalis* | MH236308, MH236312, MH236314, MH236328, MH362705, MH236319, MH236320, MH591461<sup>Δ</sup>, MH236341<sup>Δ</sup>, MH236325<sup>Δ</sup>, MH236318<sup>Δ</sup>. |
| *Staphylococcus. aureus* | OK642790<sup>◇</sup>, OK642791<sup>◇</sup>, OK642793<sup>◆</sup>, OK642796<sup>◆</sup>, OK642794, OK642795, OK642797, OK642798, OK642799, OK642792. |
| *Staphylococcus. epidermidis* | OK642800, OK642801, OK642803, OK642804. |
| *Staphylococcus. haemolyticus* | OK642805, OK642806, OK642807, OK642808, OK642809. |
| *Staphylococcus. hominis* | OK642810 |
| *Staphylococcus cohnii* | OK642811 |
| *Staphylococcus klosii* | OK642812 |
| *Staphylococcus warneri* | OK642813 |
| *Acinetobacter baumannii* | PP659668, PP659669, PP659670, PP659673,<br>PP660317, PP732465, PP660318, PP660321,<br>PP660327, PP660328, PP732466, PP660333,<br>PP660334, PP660338, PP660341, PP660342,<br>PP660550 |
| *Acinetobacter pittii* | PP660329, PP660332, PP660344, PP660545,<br>PP660551, PP660552, PP659677 |
| *Acinetobacter nosocomialis* | PP660549 |
| *Enterobacter hormaechei* | PP659673 |

△represents strains that can be lysed by Lys22 but resistant to phage LY0322. ◇ marked strains are methicillin resistant *S. aureus* (MRSA), well ◆ marked strains are resistant to both methicillin and vancomycin.

As some endolysins also showed lytic activity to Gram-negative bacteria, 44 strains of *A. baumannii*, 7 strains of *A. pittii*, 1 strain of *A. nosocomialis*, 2 strains of *E. hormaechei,* 20 strains of *Pseudomonas aeruginosa*, 21 strains of *Klebsiella pneumoniae* were used as target bacteria of Lys22. Among them, 17 strains of *A. baumannii,* all *A. pittii and A. nosocomialis*, and 1 strain of *E. hormaechei* can be lysed by Lys22 (Table. 1)^[23]^.

### Stability of Lys22

The stability of Lys22 under different conditions was investigated. When Lys22 was treated with EDTA, its killing effect remained relatively stable at concentrations ranging from 6.5 µM to 100 µM EDTA, as shown in Fig. 2A. The study also found that Lys22 keeps stable killing effects in a concentration range of 25 µM to 100 µM NaCl, as shown in Fig. 2B. This suggests that Lys22 is effective in saline environments, which is particularly relevant for its potential application in wound care. And lytic activity of Lys22 also remains stable across a wide range of pH values, from pH 4 to pH 10 (Fig. 2C). Finally, the study found that Lys22 exhibits high activity below 55°C (Fig. 2D).

**Fig. 2.**
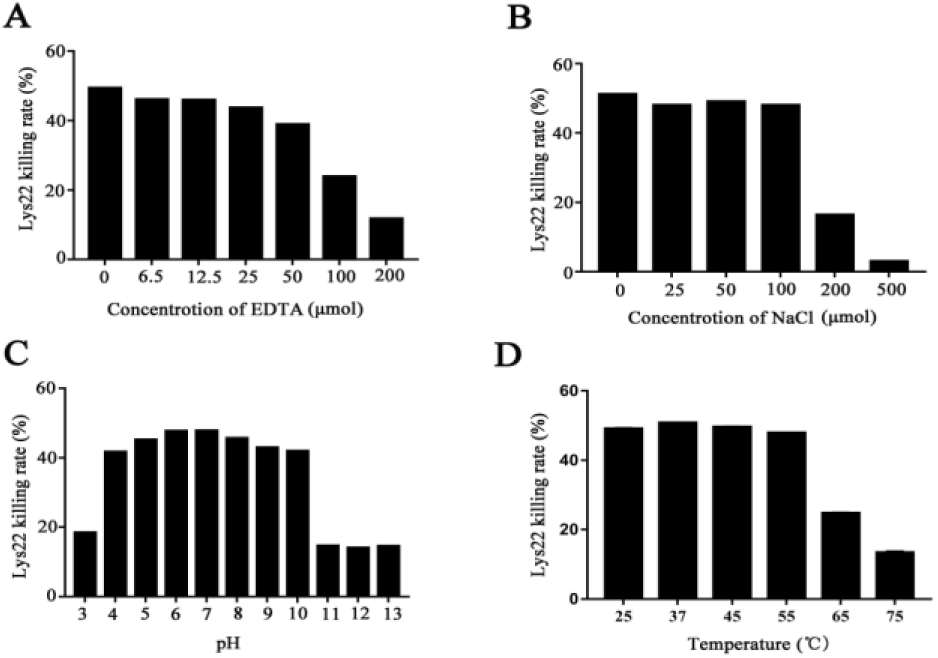
Effects of different factors on Lys22 killing activity to E. faecalis. A and B represented the effect of different concentrations of EDTA and NaCl on Lys22 killing activity. C and D represented the effect of different pH and temperature on Lys22 killing activity.

### The effect of Lys22 on *E. faecalis* biofilm

Confocal laser scanning microscopy (CLSM) was used to analyze the three-dimensional structure of *E. faecalis* biofilms with or without Lys22 (Fig. 3). The survival bacteria within the biofilm were stained by green, fluorescent dyes, while the killed bacteria were not. The biofilm formed in the control group was thicker than that in the experimental group with Lys22 whether it was added at early stage of biofilm formation or in mature biofilm. These findings suggest that Lys22 strongly inhibited *E. faecalis* biofilm formation and destroyed mature biofilm.

**Fig. 3.**
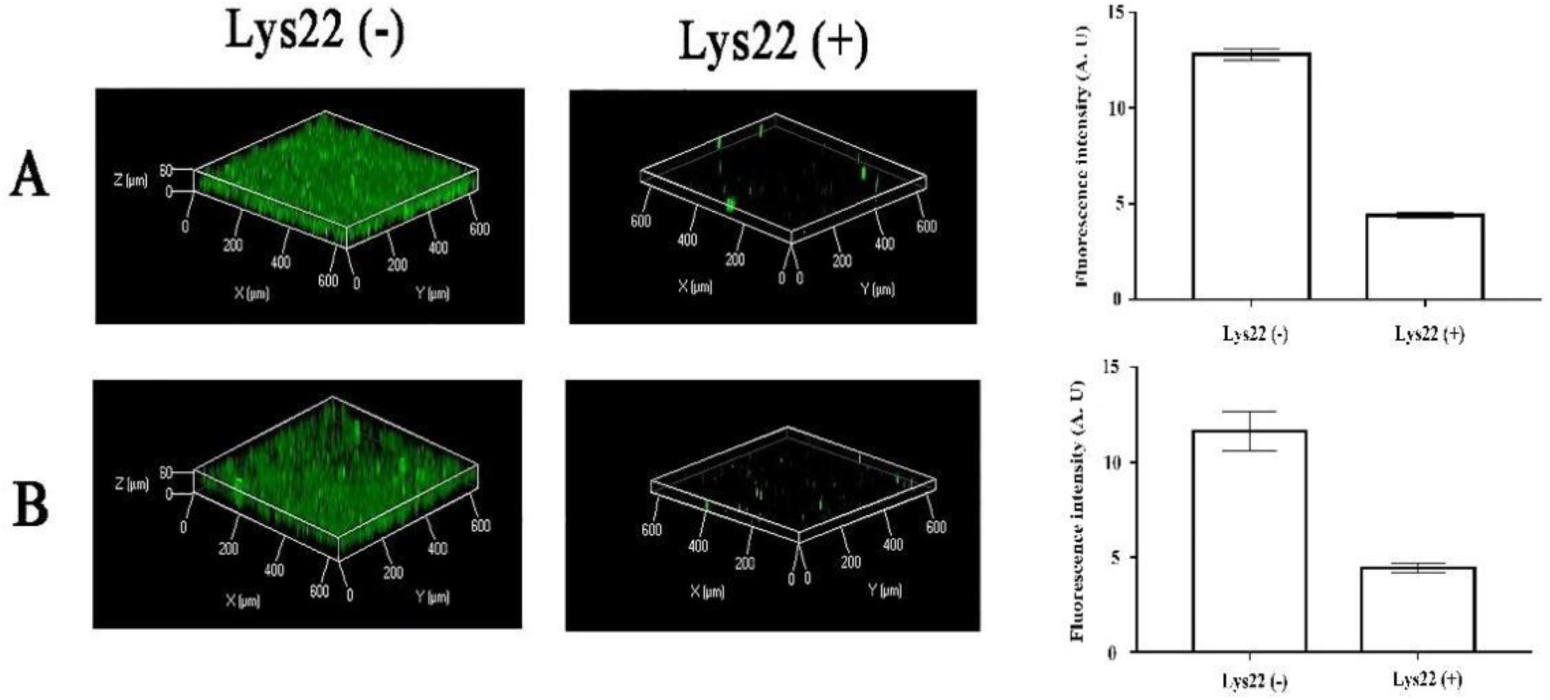
Effects of Lys22 on biofilms of *E. faecalis* under CLSM. Panel A illustrates the effects of Lys22 administration during the initial stages of biofilm formation, while Panel B demonstrates the inhibitory impact of Lys22 on mature biofilms. Surviving bacteria within the biofilm were labeled with green fluorescence, and the Z-axis represents biofilm thickness. In both Lys22-treated groups, the Z-axis heights could not be measured due to the complete disruption of the biofilms, indicating that Lys22 significantly inhibits both the formation and growth of biofilms. Additionally, the fluorescence intensity of surviving bacteria in the Lys22-treated groups was markedly reduced, further confirming the potent inhibitory effect of Lys22 on the formation and maturation of *E. faecalis* biofilms.

Then field emission scanning electron microscopy (FESEM) was used to analyze the effects of Lys22 on *E. faecalis* biofilms on dentin tablets (Fig. 4). The *E. faecalis* biofilm without Lys22 was dense and layered, the bacteria showed normal morphology and clear adhesion to neighbors. However, in the groups treated by Lys22 at early stage, the number of *E. faecalis* was significantly reduced, the biofilm could not be observed, and the survival bacteria had abnormal morphology. And in the group treated by Lys22 at the mature biofilm, except debris of lysed bacteria, no biofilm and survival bacterial could be observed.

**Fig. 4.**
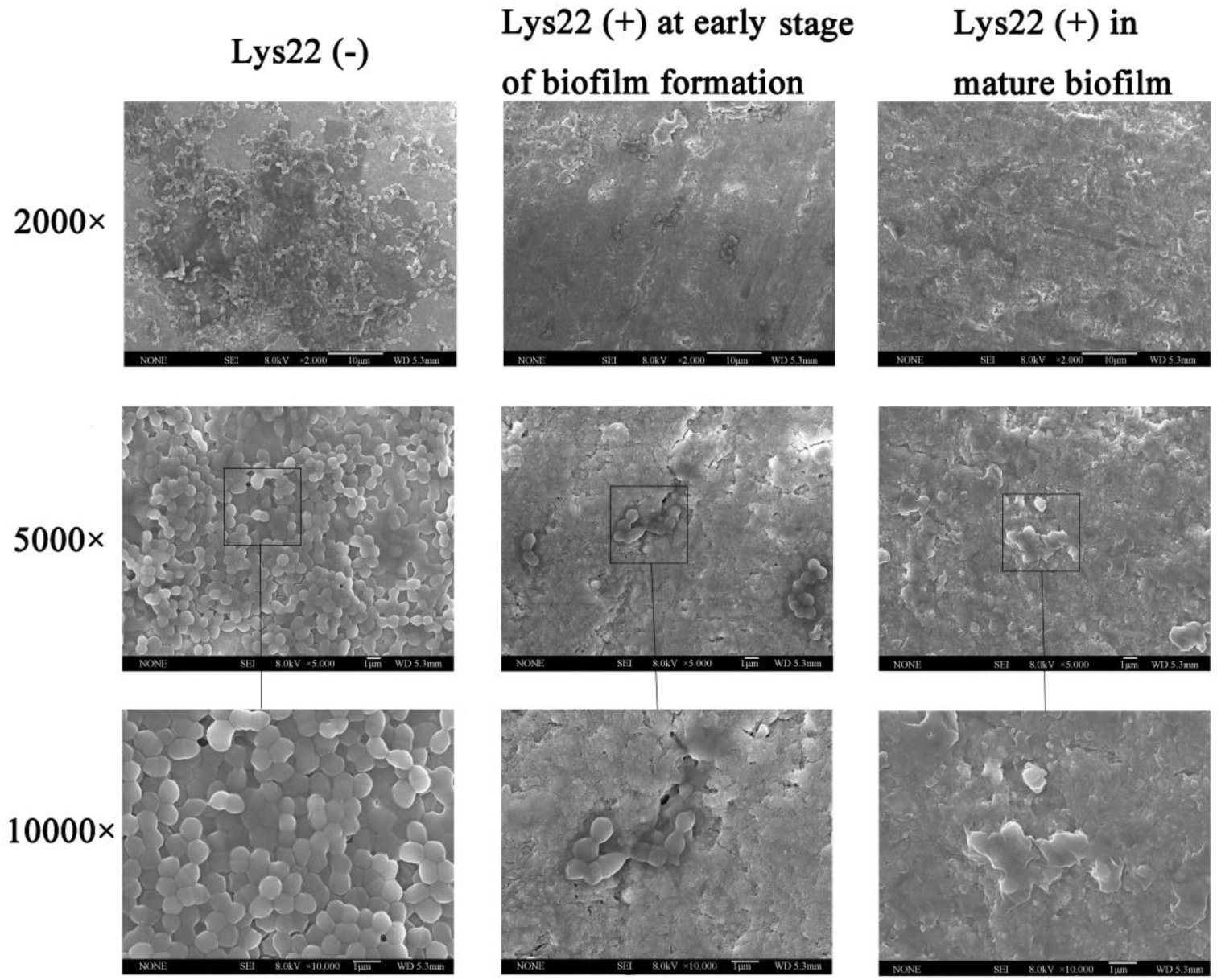
Effects of Lys22 on *E. faecalis* biofilms on dentin tablets. The groups with Lys22 treatment show the inhibition of biofilm not only at early stage but also in mature biofilm on dentin tablets. For each group, there are three images with 2000×, 5000×, and 10000× magnifications respectively.

### The effect of Lys22 against mixed biofilm

Gentamicin resistant *E. faecalis*, methicillin and vancomycin resistant *S. aureus*, and imipenem resistant *A. baumannii* were used as host bacteria to test inhibition effect of Lys22 on dual- and triple-species biofilm. The biofilm was stained by crystal violet and the optical density of all absorbed crystal violet was tested at 570 nm (Fig 5 A-D). While planktonic bacteria were evaluated by testing optical density at 600 nm (Fig. 5 E-H). The biofilm of all groups was also observed by electronic microscope (Fig. 6 and Fig.7). Compare to untreated controls, Lys22 showed strong inhibition to both dual- and triple-biofilm (*p*<0.05). The planktonic cells were also effectively inhibited at all tested stages (*p*<0.05).

**Fig. 5.**
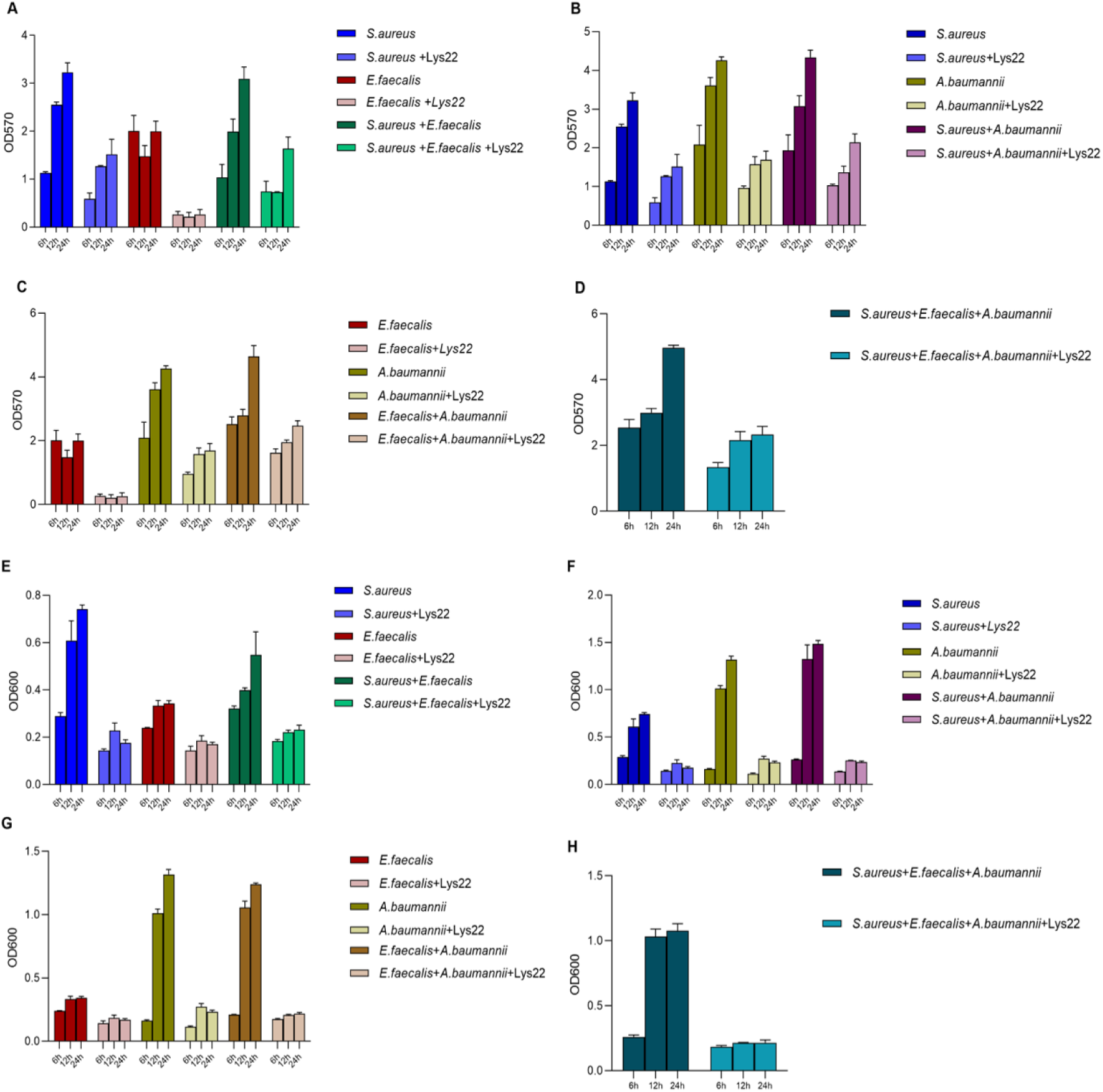
Effect of Lys22 on biofilm formation. The histograms show the impact of Lys22 treatment on the biofilm (A-D) and planktonic cell density (E-H) of *S. aureus*, *E. faecalis*, and *A. baumannii* over time points of 6, 12, and 24 hours.

**Fig. 6.**
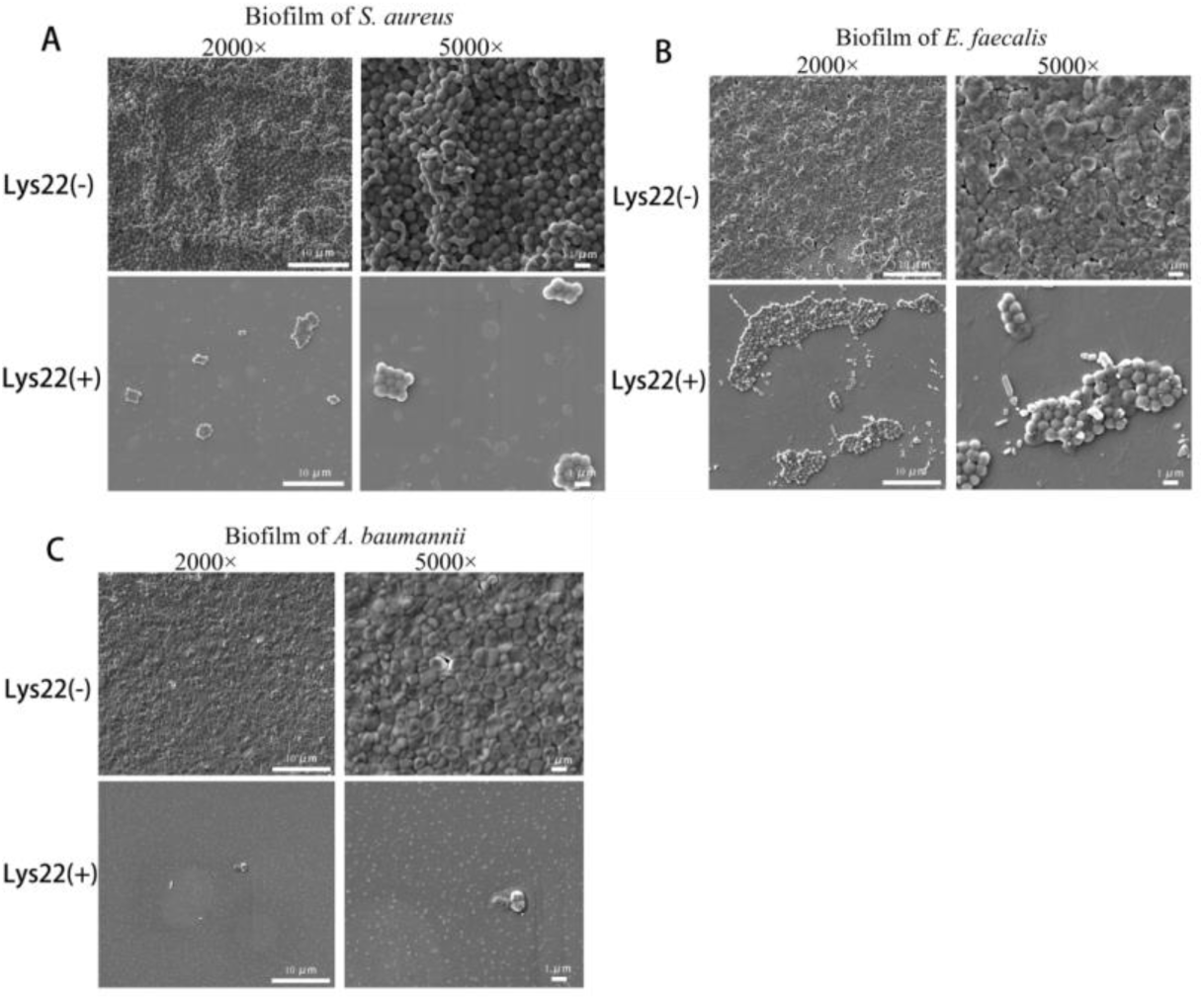
The inhibition of Lys22 to single biofilm of *S. aureus, E. faecalis* and *A. baumannii* observed under scanning electronic microscope. In all groups, the bacteria were cultured 6h. Then, Lys22 was added to the working concentration of 50 μg/mL in Lys22 treated groups. Then bacteria in all groups were cultured 3h before observed under electronic microscope. A is biofilm of *S. aureus*, B is biofilm of *E. faecalis*, and C is biofilm of *A. baumannii*.

**Fig. 7.**
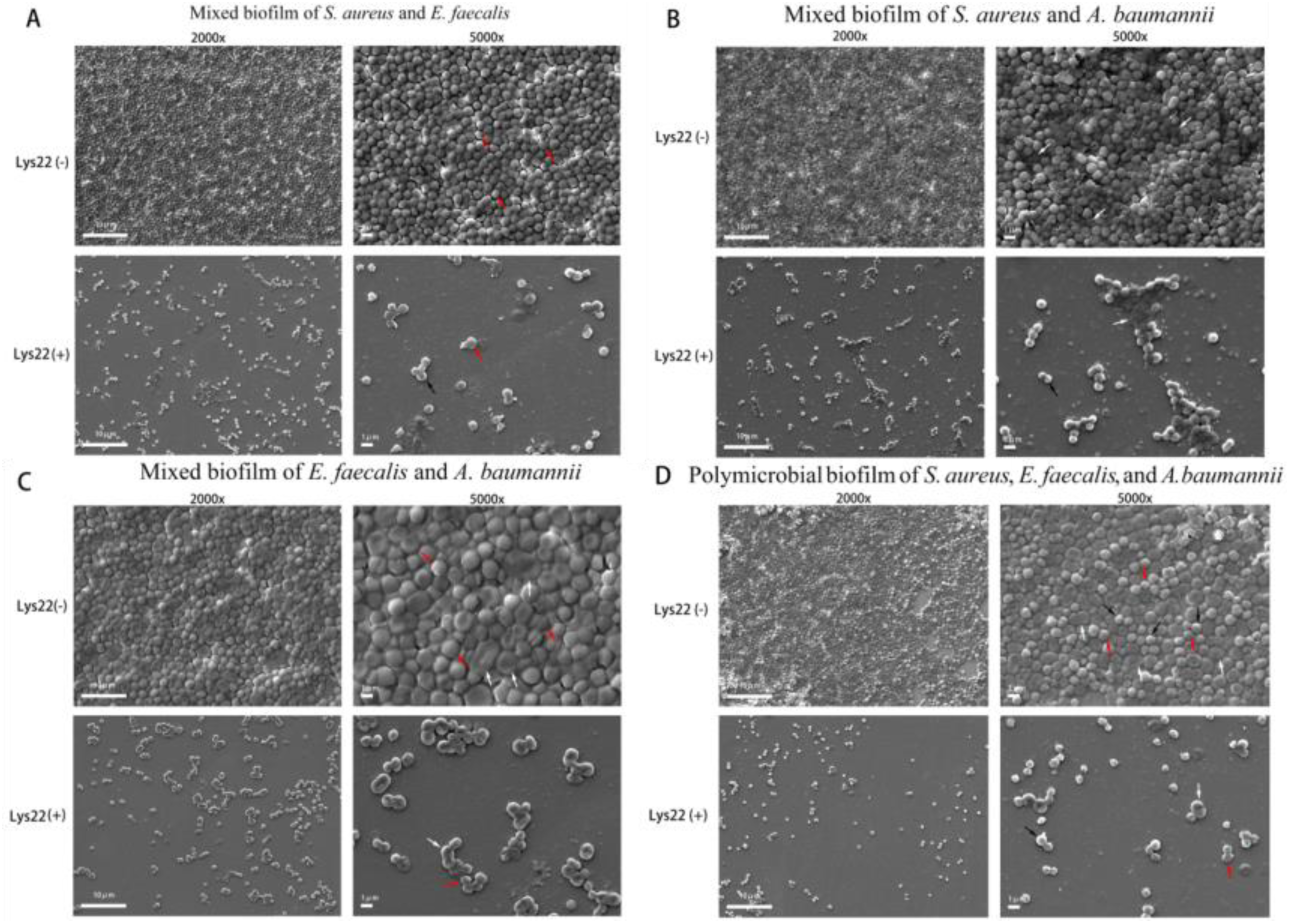
The inhibition of Lys22 to mixed biofilm of *S. aureus, E. faecalis* and *A. baumannii* observed under scanning electronic microscope. The method of this test is like that in Fig. 6. A is the mixed biofilm of *S. aureus* and *E. faecalis.* B is the mixed biofilm of *S. aureus* and *A. baumannii*. C is the mixed biofilm of *E. faecalis* and *A. baumannii*. D is the mixed biofilm of *S. aureus*, *E. faecalis*, and *A. baumannii*. Black arrows, red arrows and white arrows respectively marked *S. aureus*, *E. faecalis.* and *A. baumannii*.

### Biofilm removability by SEM

Scanning electron microscopy (SEM) observations unequivocally demonstrated that the administration of Lys22 substantially inhibited biofilm formation in all co-cultured groups, compared to the control groups where only bacterial cultures were used (Fig.6). The bacteria in the groups treated with Lys22 (50μg/mL) exhibited a marked reduction of forming continuous biofilms. Moreover, most surviving bacterial cells in these groups displayed altered morphology and arrangement compared to control groups. And the survival bacteria in the biofilm were also cultured and counted to determine the species according to the morphology of colonies (Supplementary Fig. 1).

### The effect of Lys22 on biofilm-associated gene **expression**

It has been proved that some virulent genes are related to biofilm formation and virulence of *S. aureus*, *E. faecalis*, and *A. baumannii*, and the impact of Lys22 on these genes’ expression was also tested by qRT-PCR. In *S.aureus*, the expression of key pathogenic genes, including alpha-hemolysin (*hla*, *hld*), *sarA*, and *agrA*, were decreased by Lys22 (Fig.8A)(*P*<0.05). In *E. faecalis*, plasmid-encoded aggregation factor gene (*asa-1*), cytolysin gene (*cylA*), adherence to collagen gene (*ace*), surface protein gene (*esp*), endocarditis antigen gene (*efa*), hyaluronidase gene (*hyl*) and enterococcus binding gene (*ebp*) were significantly down-regulated (Fig. 8B) (P< 0.05). And in *A. baumannii*, outer membrane protein gene (*OmpA*), lipopolysaccharide gene (*lpsB*), penicillin-binding protein gene (*pbpG*), biofilm formation regulator S gene (*bfmS*), Csu pili cluster gene’s member *csuC*, and important component of resistant island *abaR*, were downregulated by Lys22 (Fig.8C). However, *sig-B*, *icaA*, and *aur* of *S. aureus*, and *plcD* of *A. baumannii* were upregulated(*P*<0.05).

**Fig. 8.**
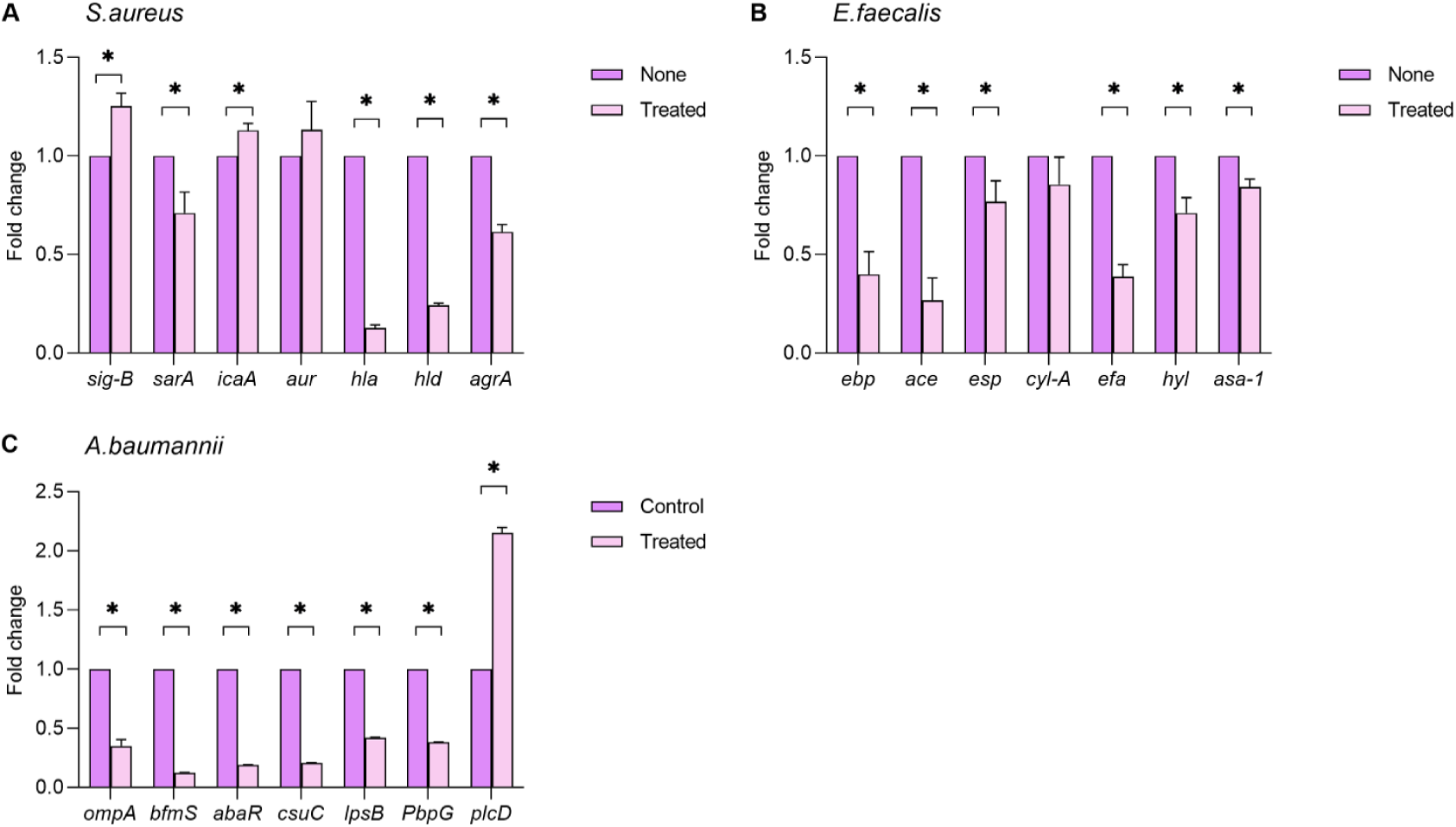
Effects of Lys22 on virulent genes transcription of *S. aureus*, *E. faecalis*, and *A. baumannii*. All bacteria were cultured in liquid medium with or without Lys22 (50 μg/mL) at 37 °C with 150 rpm shaking for 12h. Transcriptional profiles were acquired by qRT - PCR. 16SrRNA was used as a housekeeping gene. Fold changes represent a change in the transcriptions of treated vs. non - treated controls. The experiment was conducted twice and one gene was tested by three parallel qRT - PCRs per time. \**P* < 0.05 vs. non - treated controls.

### The protection of Lys22 to *S. aureus*, *E. faecalis* and *A. baumannii* attacked zebrafish embryos

The minimum lethal doses of *S. aureus*, *E. faecalis*, and *A. baumannii* to Zebra fish embryos has been experimentally validated as 1 × 10^8^ CFU/well, 1 × 10^8^ CFU/well, and 1 × 10^7^ CFU/well, respectively. The minimum lethal doses were sufficient to induce 100% mortality rate to zebra fish embryos within 24 hours. Lys22 administration dramatically increased survival rates of zebra fish embryos from lethal dose of bacterial infections (Fig. 9A) The survival rates of *S. aureus*, *E. faecalis*, and *A. baumannii* infected groups were respectively increased to 90%, 100%, and 73.33% by Lys22 treatment (*p*<0.05). Moreover, Lys22 treatment also increased the survival rates of zebra fish embryos in dual or triple species bacterial infection. The survival rates of zebra fish embryos in *S. aureus* + *E. faecalis* group, *S. aureus* + *A. baumannii* group, *E. faecalis* + *A. baumannii*, and *S. aureus* + *A. baumannii* + *E. faecalis* group were respectively increased to 100%, 60%, 70%, and 100% by Lys22 treatment (*p*< 0.05) (Fig. 9B).

**Fig. 9.**
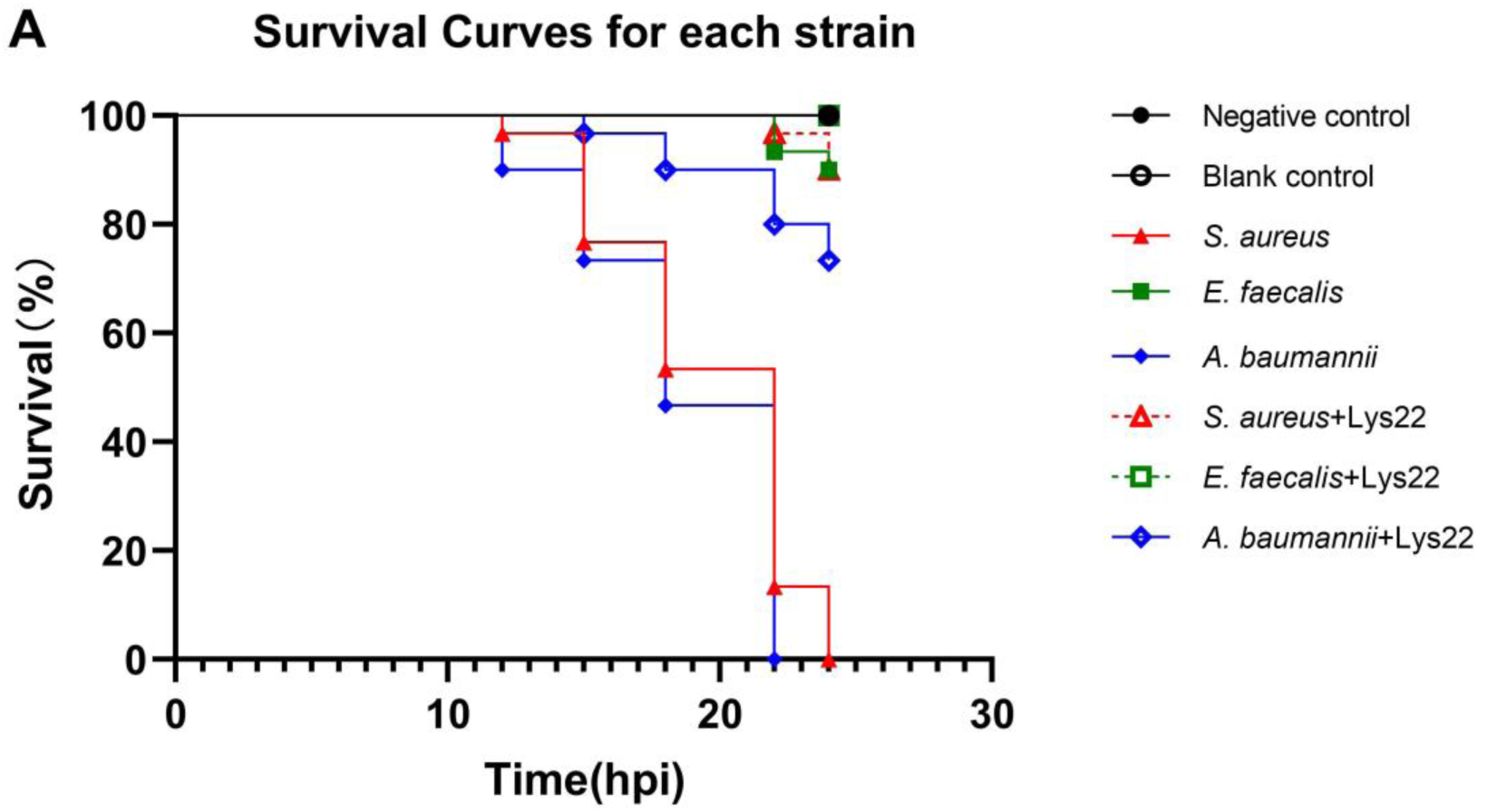
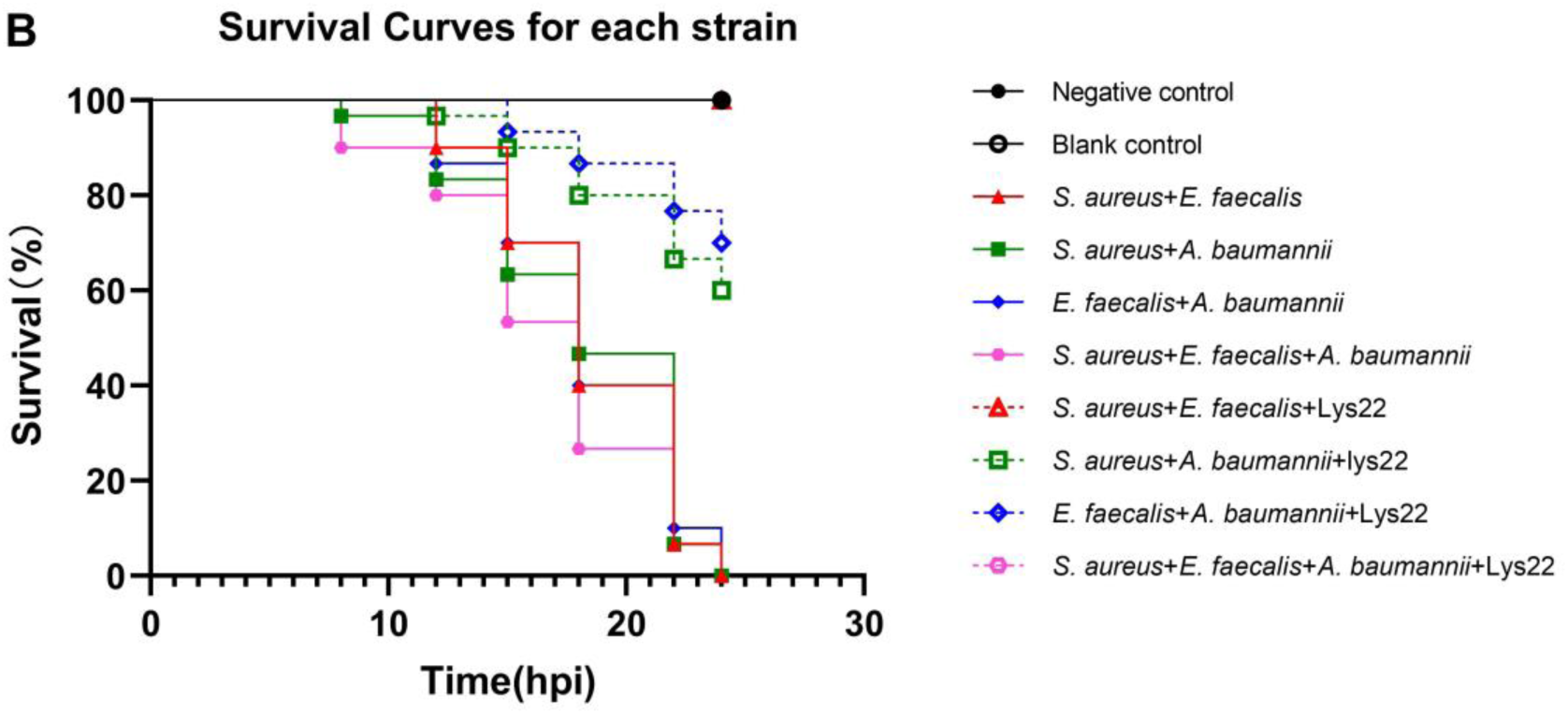
The protective effect of Lys22 on single or mixed species bacteria attacked zebrafish embryos. A is the survival curves of single species attacked group, and B is the survival curves of the groups attacked by mixed species of bacteria.

## Discussion

Endolysins are peptidoglycan hydrolases that degrade the peptidoglycan (PG) layer of the host bacterium ‘from within’ at the end of their lytic multiplication cycle. The journey of endolysin from host bacterial cytoplasm to their PG substrate needs the help of holins. Holins are also the component of phage lysis cassette of tailed phages. Holins are produced during the late stages of infection and, once a critical concentration is reached, create holes in the cytoplasmic membrane by oligomerization, allowing the endolysins, which have accumulated in the cytoplasm, to access their PG substrate ^[24]^. As Gram-positive bacteria lack outer membrane (OM), endolysins can easily reach their PG substrate and lyase cell wall when they are exogenously administrated. But the OM of Gram-negative bacteria form a barrier which prevents exogenous endolysins from accessing and degrading the underneath PG layer, thereby protecting the Gram-negative bacteria from lysin attack. So, overcoming the OM barrier of Gram-negative bacteria became very important to explore the full potential of lysins in the battle against MDR bacteria. Finding lysins with an intrinsic ability to permeate the OM, simultaneous application lysins with outer membrane permeabilizers (OMPs), and modifying the structure of lysins by engineering technique are currently proposed strategies ^[25]^.

Enterococcus phage LY0322 is a lytic phage previously isolated and identified in our lab. The sensitive host bacteria of LY0322 included 7 strains of *E. faecalis* and 2 strains of *E. faecium*. LY0322 cannot lyse any *Staphylococcus* and Gram-negative bacteria. Compared to phage LY0322, Lys22 has an outstanding wider host range, more *E. faecalis* strains, several species of *Staphylococcus* and *Acinetobacter*, and a strain of *E. hormaechei* are sensitive to the lysis of Lys22. But 2 strains of *E. faecium* sensitive to phage LY0322 were resistant to Lys22. The lytic activity to *Acinetobacter spp.* proved that Lys22 has good intrinsic permeability to OM of Gram-negative bacteria. Some strains of *Pseudomonas aeruginosa*, *Escherichia coli*, and *Klebsiella pneumonia* were also used as target bacteria in host range test, but no sensitive strains were found. And interestingly, Lys22 easily produced plaques of sensitive *Staphylococcus* and *Acinetobacter* in dropping lawn test. While the plaques in the lawn of different *E. faecalis* are usually small, nontransparent or hard to be observed. But in liquid medium, the number of survival *E. faecalis* both in biofilm and in suspension were dramatically decreased by Lys22. The wide and complicated host range of Lys22 bring us many new issues to explore and understand the mechanisms of lysin-membrane interaction and lysin-peptidoglycan target recognition.

The emergence of multidrug-resistant *S. aureus*, *E. faecalis*, and *A. baumannii* highlights the critical need for alternative therapeutic strategies. These pathogens are notorious for their ability to form robust biofilms on the surface of living or non-living organisms and medical devices, significantly hindering the effectiveness of traditional antibiotics ^[26]^. Lys22 showed an effective inhibition to both biofilm formation and mature biofilm of these three bacteria. As *E. faecalis* was detected in 4%-40% of primary root canal infections and 24%-77% of secondary root canal infections ^[27]^. Root canal therapy is a routine treatment for root canal infections, which can seal and isolate the root canal system from the source of bacterial infection. However, the failure rate of root canal therapy is as high as 10%-15% ^[28]^. The biofilm provides protection for *E. faecalis* from the attack of the environment and facilitate its survival ^[29, 30]^. Considering this factor, we tested the effect of Lys22 on biofilm of dentin slices, and found that Lys22 can inhibit the formation of biofilm and has killing effect on the bacteria in the mature biofilm. And in dual or triple bacteria mixed culture groups, the biofilms were also inhibited by Lys22.

Moreover, Lys22 induced cell wall destruction also triggered expression changes of some genes related to biofilm formation and pathogenicity. As qRT-PCR assay showed, In *S. aureus*, *hla*, *hld*, *sarA*, and *agrA* were downregulated, while *sigB* and *icaA* were upregulated (Fig.8A). *Hla* and *hld* are crucial for the pathogenicity of *S. aureus*, and *sarA* facilitate biofilm formation ^[31,32]^. And the icaA gene encodes an enzyme, N-acetylglucosamine transferase, which is involved in the synthesis of polysaccharide intercellular adhesin (PIA), a key component of the biofilm in *S. aureus* ^[33]^. Auxiliary gene regulator (*agr*) and alternative sigma factor σB (*sigB*) are regulators. *Agr* is a survival enhancing gene involved in regulating virulence and intercellular communication genes expression via two-component signaling system (TCSTS) ^[34]^. In various Gram-positive bacteria, such as *B. subtilis*, *B. cereus*, *L. monocytogenes*, and *S. aureus, sigB* is crucial for restore normal physiological function or balance when bacteria undergo stress, damage, or dysfunction ^[35]^. Upregulation of *sigB* and *icaA* suggests that *S. aureus* activates compensatory mechanisms to survive under lysin suppression.

In *E. faecalis*, *asa1*, *cylA*, *ebp*, *esp*, *efa*, and *hyl*, which involved in biofilm formation and pathogenesis were downregulated to varying degrees by Lys22 (Fig.3C). The product of *asa1* facilitates the adherence to renal tubular cells and human macrophages. The cytolysin gene (*cylA*) encodes a protein capable of lysing both prokaryotic and eukaryotic cells ^[36]^. Additionally, *ebp* and *ace* genes encoded structures are critical for biofilm formation and adhesion, which play significant roles in experimental urinary tract infections (UTIs) and endocarditis ^[37]^. The surface protein (*esp*), located in the bacterial cell wall, contributes to colonization and biofilm formation. The endocarditis antigen (*efa*) is pivotal in biofilm formation and pathogenesis ^[38]^. Hyaluronidase (*hyl*), by degrading hyaluronic acid, facilitates colonization of host tissues ^[39]^. In *A. baumannii* cells, OmpA promotes host cell apoptosis, epithelial cell adhesion, invasion, biofilm formation, surface motility, and serum resistance ^[40]^. Lipopolysaccharide (*lpsB*) helps evade the host immune response and triggers an inflammatory response ^[41]^. Penicillin-binding protein (*pbpG*) plays a role in peptidoglycan biosynthesis, cell stability, and serum growth ^[42]^. The *abaR* islands are recognized as potential contributors to antibiotic resistance and heavy metal resistance ^[43]^.

In *A. baumannii*, virulent genes involved in biofilm forming *(csuC, BfmS, plcD*), iron acquisition (*IpsB*), encoding outer membrane protein (*ompA*), surface glycoconjugates (PbpG), secretory system (*ompA*, *plcD*), and quorum sensing system (*abaR*) were tested in our study ^[44,45,46,47,48]^. All tested genes other than *plcD* were down regulated by Lys22. *PlcD* is very important to *A. baumannii* survival because the product of *PlcD* is implicated in biofilm formation, serum resistance and antibiotic resistance. Like that of *S. aureus*, a compensatory activation of protective system may play a role under lysin suppression.

The Lys22 treatment significantly protected zebrafish from lethal infections caused by various bacterial species, including combinations of *E. faecalis, S. aureus,* and *A. baumannii.* Lys22 effectively protected zebrafish from MASA infections, with near-complete protection observed. However, its efficacy was reduced against the Gram-negative bacterium *A. baumannii*, potentially due to the presence of bacterial metabolites, resembling lipid A (endotoxin) ^[49]^, which may have contributed to host mortality prior to the action of the endolysin. Interestingly, when the three bacteria were co-infected, Lys22 conferred full protection, likely due to inter-bacterial competition in the mixed culture, which reduced the number of pathogenic strains capable of causing zebrafish mortality. Overall, Lys22 demonstrated significant therapeutic potential in *in vivo* infection models.

## Importance

This study confirms that endolysin Lys22 has a wide host range including *E. faecalis*, *Staphylococcus spp.* and *Acinetobacter spp.* in disrupting biofilms and combating multidrug-resistant bacterial infections. The lytic activity to Acinetobacter spp. proved Lys22 has good intrinsic permeability to OM of Gram-negative bacteria. Lys22’s ability to destroy single-species biofilm and mixed biofilm of *E. faecalis, S. aureus* and *A. baumannii* offers a distinct advantage over traditional antibiotics, especially in overcoming polymicrobial biofilm resistance. And Lys22 effectively protected zebrafish eggs attacked by single, dual and triple species attacks of *E. faecalis, S. aureus* and *A. baumannii.* Additionally, Lys22 effectively downregulated virulence genes such as *agrA* and *icaA* in *S. aureus* and *asa1, cylA,* and *hyl* in *E. faecalis*, *OmpA* and *lpsB* in *A. baumannii*, reducing their pathogenicity.

## Methods

### Bacterial strains and zebrafish embryos

All bacteria used in this study are isolated from Affiliated Hospital of Changchun University of Chinese Medicine and identified by Matrix Assisted Laser Desorption Ionization (MALDI)-Time of Light (TOF)-Mass Spectrometer (MS) after purification. Then 16SrDNA were multiplied and analyzed by routine methods. All 16S rDNA sequences were submitted to NCBI and accession numbers were listed in Table 1. The minimum inhibitory concentration (MIC) of target bacteria were tested according Clinical and Laboratory Standards Institute (CLSI)1 method (http://em100.edaptivedocs.net/dashboard.aspx.).

Unhatched zebrafish embryos (0-48hpf) used in this study are from AB strains.

### The expression of Lys22

Gene ID of Lys22 in NCBI is 40101561. Gene of Lys22 was cloned into pET28a (+) vector with 6×His label according to universal method ^[50]^. The plasmid was transformed into *E. coli* BL21 by heat shock method. After successful transformation, E. coli BL21 was incubated in LB medium containing 0.1 mg/mL of kanamycin and cultured at 37℃until OD600 reached between 0.4 to 0.6. Then Lys22 expression was induced by 0.1 mM isopropyl β-D-thiogalactoside (IPTG) at 20 ℃ for 16 h. The supernatant of culture medium was collected by centrifugation at 4℃, 8000 g for 10 min. To remove miscellaneous proteins and a part of endotoxins, the supernatant was filtrated with a 100 KDa ultrafiltration tube (Milipore) at 4 ℃ and 5000 g for 15 min. The obtained filtrate was further concentrated with a 10 KDa ultrafiltration tube at 4℃ and 5000 g centrifugation for 15 min. The concentrated proteins were washed and collected with Lysis Buffer [50 mmol Tris-HCl (pH 8.0)] and analyzed by SDS-PAGE. Finally, His-tagged Lys22 were purified by Ni-NTA and collected by 40 mM to 250 mM imidazole washing. Concentrations of Lys22 in series of imidazole elution was determined by BCA protein determination kit ^[51]^. Amino acid sequences of expressed Lys22 were performed to analyze their Zn+ binding sites using the SWISS-MODEL server ^[52]^.

### Determination of the host range of endolysin Lys22

The frozen tubes with bacteria were placed in 37°C water immediately after being taken out of -80 °C refrigerator. After rapid thaw, 100μL stored bacteria were inoculated into 1 mL liquid BHI and cultured in shaking incubator at 37°C, 150 rpm for 1 h. Then all bacteria were inoculated regionally at BHI solid medium and cultured overnight to get purified colony at 37 °C. Subsequently, single colonies were selected and inoculated into LB broth and cultured at 37°C with shaking at 150 rpm until reaching the logarithmic phase (OD_600_ = 0.5-0.6). Then, 100 μL of bacterial culture was evenly inoculated to use a sterile spreader to evenly cover LB agar plates. After the surface became dry, 5 μL of endolysin Lys22 was dropped to the surface of the agar, and the plates were incubated overnight at 37°C. The presence of lysis zones was observed to determine the susceptibility of the strains to Lys22. Lysis zones indicate effective lysis by Lys22, while their absence suggests that the strains are resistant to the lysin ^[23]^.

### The stability of Lys22

To evaluate the viability as a potential therapeutic agent, the stability of Lys22 under different conditions was tested according to universal methods with small modification ^[53,54]^. *E. faecalis*, with 16S rRNA accession number OK642793, was used in these experiments.

To test the lytic activity of Lys22 under different concentration of NaCl or EDTA, freshly cultured *E. faecalis* was suspended in phosphate buffer saline (PBS, pH 7.4) to a final OD600 of 0.5, containing various concentrations of EDTA ranging from 0 to 800 µM or NaCl ranging from 0 to 1000 µM. Then, Lys22 was added into the mixture and incubated 2h at 37°C. Finally, the OD600 of the mixtures were tested to evaluate the lytic activity of Lys22. Similarly, the lytic activities of Lys22 at different pH values were also measured using a universal method.

To evaluate the thermal stability of Lys22, Lys22 was treated at different temperature including 25°C, 37°C, 45°C, 55°C, 65°C and 75°C for 30 min. After treatment, Lys22 was added into PBS suspended *E. faecalis*. The killing effect of Lys22 against *E. faecalis* was tested by measuring the decrease in OD600 within 100 min ^[54]^.

In all above tests, the final concentration of Lys22 was adjusted to 50 µg/ mL according MIC tests (supplement documents).

## The effect of Lys22 on *E. faecalis* biofilm

### Confocal Laser Scanning Microscopy (CLSM) analysis

In 24 well plates, Lys22 was added into the *E. faecalis* suspension to achieve a median concentration of 50 μg/mL. After 6 h of incubation, the biofilm in each well was washed with 1 mL 0.01M PBS and stained by Bacterial Viability Kit (Molecular Probes) according to attached procedures. Following another washing step with 0.01M PBS, the stained biofilm was air-dried for 10 minutes and fixed with 1 mL of 2.5% glutaraldehyde (Solarbio, Beijing Co, Ltd.) for 30 minutes at room temperature. Finally, we washed the biofilm again with 0.01M PBS and analyzed it using a Leica Ultra View VOX confocal laser scanning microscope (Leica Microsystems), as described in a previous paper ^[55]^

### Inhibition of biofilm in human dentin slices by Lys22

Briefly, the fully developed teeth were obtained from Stomatology Hospital of Jilin University. The crown was removed using a gold needle, after which the dentin was cut into sections measuring 5 mm × 5 mm × 1 mm. Once the dentin sections were obtained, they were autoclaved at 121°C for 15 minutes, as described in a previous paper ^[56]^

To study the impact of Lys22 on the inhibition of biofilm formation, *E. faecalis* was inoculated onto sterilized detin slices in the presence or absence of Lys22 at a concentration of 50 μg/mL1, and then cultured for 6h at 37°C in a 24 well plate. After incubation, the dentin slices were washed with 0.01M PBS three times, air-dried at room temperature, and fixed with 2.5% glutaraldehyde for 30 minutes. Then, the buffer was replaced with increasing concentrations of ethanol (70%, 80%, 90%, 95%, and 100% for 15 minutes each) to dehydrate the slices. Finally, the dried slices were sprayed with gold and observed under a field emission scanning electron microscope (FESEM, JSM-6700F, JEOL, Japan, 8.0KeV), as described in previous papers ^[53]^.

To study the effect of Lys22 on mature biofilm, *E. faecalis* was inoculated onto sterilized dentin slices and cultured for 6h at 37°C in a 24-well plate firstly. Next, the plate was washed with 0.01M PBS and the medium in experimental wells was replaced with 1 mL of BHI medium containing Lys22 at a concentration of 50 μg/mL. The dentin slice was then treated and observed using the same method as described above.

### The effect of Lys22 on mixed biofilm

#### Turbidity test and crystal violet method

Except *E. faecalis* used in upper experiments, Methicillin and vancomycin resistant *S. aureus* and imipenem resistant *A. baumannii* were selected as host bacteria in this experiment and subsequent experiments. 16S rDNA accession nmubers of *S. aureus*, and *A. baumannii* are PP660341 and MH236318 respectively. All bacteria were overnight cultured in BHI solid medium at 37°C. The single colonies were inoculated into BHI broth and cultured at 37°C with shaking at 150 rpm for 6 hours until reaching the logarithmic phase (OD_600_=0.5-0.6). Then concentrated Lys22 (500 μg/mL) was mixed into bacterial cultures and adjusted its working concentration to 50 μg/mL. Subsequently, the mixture was added into 96-well plates at 200 μL per well and incubated at 37°C. The planktonic bacteria and biofilm were tested at 6, 12, and 24 hours. For planktonic bacteria evaluation, the culture media were gently absorbed out and added into new 96-well plates to test OD_600_ (BioTek, USA). For biofilm evaluation, the plates were washed three times with 200 μL of 0.1 M PBS and air-dried at room temperature after culture media being removed. Subsequently, 200 μL of 0.1% crystal violet was added into each well and kept for 10 minutes. Then the solutions were taken out, and the wells were gently washed three times with 0.1 M PBS and air-dried at room temperature. Finally, 200 μL of 95% ethanol was added to each well and maintained over 10 minutes to release crystal violet combined in biofilms. And the OD_570_ of released crystal violet was tested by microplate spectrophotometer to quantify biofilm content ^[57]^. The experiments were repeated three times with at least three parallel wells in each group.

#### Scanning electronic microscope observation

*S. aureus*, *E. faecalis* and *A. baumannii* were cultured to logarithmic phase (OD_600_=0.5-0.6) in 5 mL of LB or BHI medium at 37°C and 150 rpm. The concentration of Lys22 was adjusted to 100 μg/mL by liquid LB and BHI medium. The cultures were centrifuged, resuspended in an equal volume of the respective medium, and then 500 μL of each bacterial culture was mixed with an equal volume of 100 μg/mL Lys22. This mixture was added to each well of a 24-well plate, ensuring an initial inoculum of 10^8^ CFU for each bacterium; the wells were covered with a sterile glass sheet of 14 mm diameter. For controls, only the bacterial culture was added. The plates were incubated after 3h, non-adherent cells were removed by aspiration and the wells were washed thrice with sterile PBS. The biofilms were then freeze-dried and examined using scanning electron microscopy (SEM) (JEOL Ltd., JSM-7900F, 1-2 Musashino 3-chome, Akishima, Tokyo 196-8558, Japan.) ^[58]^.

### Quantitative Real-Time PCR (qRT-PCR)

*S. aureus*, *E. faecalis*, and *A. baumannii* were inoculated into 50 mL of LB broth and incubated at 37°C with shaking at 150 rpm for 6h and adjusted the bacteria concentration to 10^8^ CFU/mL. Subsequently, Lys22 were added into cultures at working concentration of 50 μg/mL. The control cultures were added same volume of ddH_2_O. Then, the cultures were cultured 6h again at same condition. To prevent RNA degradation, RNase inhibitor (Thermo Fisher Scientific, USA.) was added, and cells were immediately cooled in a dry ice-ethanol bath (95% ethanol) for 30 seconds. Cells were then harvested by centrifugation at 12,000 rpm for 1 minute at 4°C, and total RNA was isolated using the Total RNA Kit (Yisheng Biotechnology, China). Real-time quantitative reverse transcription polymerase chain reaction (qRT-PCR) was employed to analyze the transcript levels of virulence genes, including *agrA*, *aur*, *hla*, *hld*, *icaA*, *sarA*, and *sigB* in *S. aureus*; *efa*, *ace*, *esp*, *ebp*, *cylA*, *hyl*, and *asa1* in *E. faecalis*; and *ompA*, *bfmS*, *abaR*, *csuC*, *lpsB*, *PbpG*, and *plcD* in *A. baumannii* (Supplementary Table 1)^41^. qRT-PCR reactions were performed using SYBR Green Premix (Yaenzyme Biotechnology, China) and a real-time fluorescent quantitative PCR system (Thermo Fisher Scientific, USA). Gene-specific primers were used, and 16s rRNA served as a housekeeping gene (Supplementary Table 1) to standardize the quantification of target gene expression ^[59]^.

### Protection of Lys22 to Zebrafish embryos

*The single colony of Enterococcus faecalis*, *Staphylococcus aureus*, and *Acinetobacter baumannii* were streaked out and cultured in LB or BHI broth at 37°C, 120 rpm to Logarithmic period (OD600≈0.6). Then bacteria were centrifuged for 10 min at 9000 rpm (Hettich Universal 320/320R centrifuge) and resuspended in zebrafish embryos culture medium (Fishbio-008). Then the bacteria solution was series diluted to 10^6^, 10^7^, 10^8^, and 10^9^ CFU/mL by PBS. Zebrafish-specific culture medium and unhatched zebrafish embryos were added into 6 well plates with 6 mL medium and 30 embryos per well. Then, different volumes of prepared bacteria solution were added into 6 well plates. Subsequently, all plates were cultured at 28°C for 24h to test minimal lethal dose (MLD) of each strain. Once the MLD had been determined, this concentration of bacteria was used as the infective inoculum (challenge dose). To investigate the protection effect of Lys22, lys22 was added into each well of zebrafish embryos culture plates shortly before performing the MLD of *Enterococcus faecalis*, *Staphylococcus aureus*, and *Acinetobacter baumannii*. The living zebrafish embryos were observed and calculated at 4h, 8h, 12h, 15h, 18h, 22h, and 24h after bacteria challenge. To ensure the death of zebrafish embryos were induced by bacteria attack, a blank control without pathogens nor endolysin Lys22 and a negative control with endolysin Lys22 only were also performed at same time ^[60-62]^.

### Statistical analysis

The number of replicates for the assays is provided above, with results presented as means ± standard deviations. Statistical analysis was carried out using one-way ANOVA followed by Dunnett’s test, utilizing SPSS version 23 (SPSS Inc., Chicago, IL, USA). A *P* value of < 0.05 was considered significant, and asterisks denote significant differences between treated and untreated groups.

---

**Table 1.** The host range of Lys22.

| Strain number | Bacterial strain | Sensitivity to Lys22 |
| --- | --- | --- |
| 1 | *A. baumannii*(A1) | ++ |
| 2 | *A. baumannii*(A2) | ++ |
| 3 | *A. baumannii*(A3) | ++ |
| 4 | *A. baumannii*(A4) | - |
| 5 | *A. baumannii*(A5) | - |
| 6 | *Enterobacter hormaechei* | + |
| 7 | *A. baumannii*(A7) | - |
| 8 | *A. baumannii*(A8) | - |
| 9 | *A. baumannii*(A9) | - |
| 10 | *A. pittii*(A10) | ++ |
| 11 | *A. baumannii*(A11) | - |
| 12 | *A. baumannii*(A12) | - |
| 13 | *A. baumannii*(A13) | - |
| 14 | *A. baumannii*(A14) | - |
| 15 | *A. baumannii*(A15) | - |
| 16 | *A. baumannii*(A16) | ++ |
| 17 | *A. baumannii*(A17) | ++ |
| 18 | *A. baumannii*(A18) | ++ |
| 19 | *Enterobacter hormaechei* | - |
| 20 | *A. baumannii*(A20) | - |
| 21 | *A. baumannii*(A21) | + |
| 22 | *A. baumannii*(A22) | - |
| 23 | *A. baumannii*(A23) | - |
| 24 | *A. baumannii*(A24) | - |
| 25 | *A. baumannii*(A25) | - |
| 26 | *A. baumannii*(A26) | - |
| 27 | *A. baumannii*(A27) | ++ |
| 28 | *A. baumannii*(A28) | ++ |
| 29 | *A. baumannii*(A29) | - |
| 30 | *A. pittii*(A30) | ++ |
| 31 | *A. pittii*(A31) | ++ |
| 32 | *A. baumannii*(A32) | ++ |
| 33 | *A. baumannii*(A33) | ++ |
| 34 | *A. baumannii*(A34) | - |
| 35 | *A. baumannii*(A35) | - |
| 36 | *A. baumannii*(A36) | - |
| 37 | *A. baumannii*(A37) | ++ |
| 38 | *A. baumannii*(A38) | - |
| 39 | *A. baumannii*(A39) | - |
| 40 | *A. baumannii*(A40) | ++ |
| 41 | *A. baumannii*(A41) | ++ |
| 42 | *A. baumannii*(A42) | - |
| 43 | *A. pittii*(A43) | ++ |
| 44 | *A. baumannii*(A44) | - |
| 45 | *A. baumannii*(A45) | - |
| 46 | *A. pittii*(A46) | ++ |
| 47 | *A. baumannii*(A47) | - |
| 48 | *A. baumannii*(A48) | - |
| 49 | *A. baumannii*(A49) | - |
| 50 | *A. Nosocomialis*(A50) | ++ |
| 51 | *A. baumannii*(A51) | ++ |
| 52 | *A. pittii*(A52) | ++ |
| 53 | *A. pittii*(A53) | ++ |
| 54 | *A. baumannii*(A54) | - |

Note: ++:Highly sensitive +: Sensitive - :Nonsensitive

